# Interactions between molecular chaperone P20 and Cyt2Ba7 toxin in *Bacillus thuringiensis*

**DOI:** 10.1101/129833

**Authors:** Yongxia Shi, Mujin Tang, Yalin Liao, Wei Xu

## Abstract

P20 or 20-kilodalton protein is a molecular chaperone protein in *Bacillus thuringiensis* (*Bt*) which can increase yields and facilitates crystal formation of various insecticidal crystal proteins (ICPs). In previous studies, a *B. thuringiensis* insecticidal protein gene, *cyt*2Ba7, was cloned, expressed but its expression level is very low in *B. thuringiensis*. In this study, various expression vectors were constructed by incorporating *p*20 in forward or reverse direction in the upstream of *cyt*2Ba7 and transformed into a *B. thuringiensis* acrystalliferous strain 4Q7. The result showed that in the presence of P20, the expression of Cyt2Ba7 was significantly increased. Especially when *p*20 gene was reversely inserted in the upstream of *cyt*2Ba7 gene, the expression of Cyt2Ba7 was increased ∼3.2 times meanwhile more and bigger crystals were observed under electron microscopy. By using purified Cyt2Ba7, P20 protein and P20-specific antiserum, immunoblotting and ligand blot analysis demonstrated a strong binding affinity between P20 and Cyt2Ba7. These results reveal that P20 can promote the crystal formation and enhance the expression of Cyt2Ba7 as a molecular chaperone, which can be a powerful tool to boost the ICPs production in *B. thuringiensis* and help develop more effective insect control strategies.

## Introduction

*Bacillus thuringiensis* (*Bt*) is a gram-positive soil bacterium which can produce insecticidal crystal proteins (ICPs) that are composed of one or several toxic proteins with insecticidal and cytolytic activities [Crickmore et al., 1998]. Cry and Cyt are two major ICP groups but they do not share sequence homologies [Crickmore et al., 1998; Palma et al., 2014; Rajamohan et al., 1998]. Cry proteins are not only toxic to the larvae of lepidopteran, dipteran and coleopteran insects, but also to nematodes, protozoan pathogens, animal-parasitic liver flukes and mites [Crickmore et al., 1998; Zhang et al., 2016]. Cyt proteins are primarily lethal to dipteran insects and cytolytic to a broad range of cells including erythrocytes [Crickmore et al., 1998; Thomas and Ellar, 1983]. Its cytolytic properties are attributed to its affinity for unsaturated fatty acids in cell membranes, in which it apparently aggregates, leading to the formation of pores that cause cell lysis [Canton et al., 2014; Guerchicoff et al., 1997; Koni and Ellar, 1993, 1994; Zhang et al., 2016]. Cyt proteins appear to have a different mode of action and lack significant cross-resistance to Cry proteins, indicating that Cyt proteins may play an even more important long-term role in managing resistance to Cry proteins [Canton et al., 2014; Sayyed et al., 2001; Wirth et al., 2000; Wu and Federici, 1993; Yu et al., 2002b].

A *cyt*2 (*cyt*2Ba7) gene from a soil-isolated *B. thuringiensis* strain was cloned, expressed in *Escherichia coli* and purified to obtain the antiserum from rabbits [Yu et al., 2002b]. Purified Cyt2Ba7 also showed cytolytic activity to insect cells and mosquito larvicidal activity. However, the expression of Cyt2Ba7 in *B. thuringiensis* is very low and can only be detected by specific antiserum [Yu et al., 2002b]. Improving the synthesis of ICP proteins in *B. thuringiensis* is critical since higher yields leads to higher insecticidal activity, which could be achieved by incorporating genetic elements to regulate protein synthesis at the transcriptional, translational and posttranslational levels [Adams et al., 1989; Diaz-Mendoza et al., 2012; Elleuch et al., 2016a; Elleuch et al., 2016b; Elleuch et al., 2015; Park et al., 2007; Wu and Federici, 1993].

Molecular chaperones are proteins that help in the folding/unfolding or the assembly/disassembly of other macromolecules to form new structures or complexes. However, they do not exist in the newly formed macromolecules and function [Hartl et al., 2011]. Therefore, molecular chaperones play a critical role in the conformational quality control of the proteome by interacting with, stabilizing and remodeling a wide range of polypeptides [Hartl et al., 2011]. A group of molecular chaperones including P20 [Wu and Federici, 1993], P19 [Manasherob et al., 2001], Orf1 and Orf2 [Tang et al., 2003], have been found and characterized in the synthesis of crystal proteins in *B. thuringiensis*, in which P20 is the mostly studied molecular chaperone in *B. thuringiensis*. It is an accessory protein in *B. thuringiensis* subsp. *Israelensis* (*Bti*), which was first described during a study of Cyt1A expression [Adams et al., 1989; Wu and Federici, 1993]. Investigations concentrated on the role of P20 in Cyt1A expression proved that it is necessary for Cyt1A crystallization and host cell viability [Adams et al., 1989; Wu and Federici, 1993]. Moreover, P20 also increased the production of other *B. thuringiensis* toxins [Elleuch et al., 2016a; Elleuch et al., 2016b; Elleuch et al., 2015; Nisnevitch et al., 2006; Wang et al., 1997; Wu and Federici, 1995; Xu et al., 2001], which are poorly expressed without P20. An increase in their production in the presence of P20 prompted us to investigate if and how the 20-kDa chaperone enhances synthesis and crystallization of the Cyt2Ba7.

In this study, *p*20 and *cyt*2Ba7 were incorporated in the same vectors and transformed to an acrystalliferous *B. thuringiensis* strain. Interestingly, Cyt2Ba7 production is increased and bigger crystals were formed in the presence of P20. Then we further applied binding assay study the protein-protein interactions between P20 and Cyt2Ba7.

## Experimental Procedures

### Bacterial strains, antiserum, media and plasmids

Bacterial strains and plasmids used in this study are all deposited in State Key Laboratory for Biocontrol (Sun Yat-sen University, China). *B. thurigiensis* subsp. strain 4Q7, a plasmid-free derivate of *B. thuringiensis* serotype israelensis (*Bti*), was obtained from *Bacillus* Genetics Stock Center, Colombus, Ohio. *E. coli* strain TG1 was grown in Luria-Bertani (LB) broth or agar plate at 37°C for plasmid propagation. Plasmids pHT3101, pMD18-T, pHT301 [Yu et al., 2002b] which contains the full-length *cyt*2Ba7 gene with promoter, and pUP20 [Shi et al., 2006] which contains *cry*1Ac promoter and *p*20 gene were used for plasmid construction. Bacteria were grown in LB medium at 37°C or G-tris medium at 30°C with ampicillin (100 μg/ml) or erythromycin (25 μg/ml). Cyt2Ba7-specific antiserum, purified P20 protein and P20-specific antiserum were prepared as described [Tang, 2003; Yu et al., 2002b]. All restriction enzymes and ligation enzymes were purchased from Roche (Germany).

### Plasmids Construction

The full-length *cyt*2Ba7 gene containing native promoter and ORF sequences were amplified from plasmid pHT301 [Yu et al., 2002b] by gene-specific primers; forward primer 5’-<u>GTCGAC</u>GATAATGAGGTTATTTTGT-3’ (*Sal*I) and reverse primer 5’-<u>GCATGC</u>ATCTTACGATTTTATTGGAT-3’ (*Sph*I) as follows: 95 °C for 3 min; 35 cycles at 95 °C for 30 s, 50 °C for 30 s and 72 °C for 45 s; and final extension at 72 °C for 10 min. PCR products were ligated to vector pMD18-T and the ligation products were transformed into *E. coli* strain TG1 for DNA sequencing. The correct plasmid was digested by *Sal*I/*Sph*I and *cyt*2Ba7 fragment was subsequently ligated into vector pUP20 [Shi et al., 2006] to construct plasmid pUP20CT (Fig. 1A). Then the fragment containing a *cry*1Ac promoter, a *p*20 gene and a *cyt*2Ba7 full length gene was digested by *Bam*HI/*Sph*I from pUP20CT and ligated to pHT3101 to obtain plasmid pHP20CT, which contains a *cry*1Ac promoter and a *p*20 gene sequences upstream of full *cyt*2Ba7 gene in the same transcription direction (Fig. 1A and B). Similarly, the *cry*1Ac promoter and *p*20 gene were digested from the plasmid pUP20C by *Kpn*I/*Sph*I and then ligated into pHT301 [Yu et al., 2002b] to get plasmid pHP20CRT, which contains an upstream *cry*1Ac promoter and *p*20 gene in a reverse transcription direction to *cyt*2Ba7 gene (Fig. 1A and B). The plasmids pHT3101 (negative control), pHT301, pHP20CT and pHP20CRT were further electroporated into the acrystalliferous strain 4Q7.

**Fig. 1.**
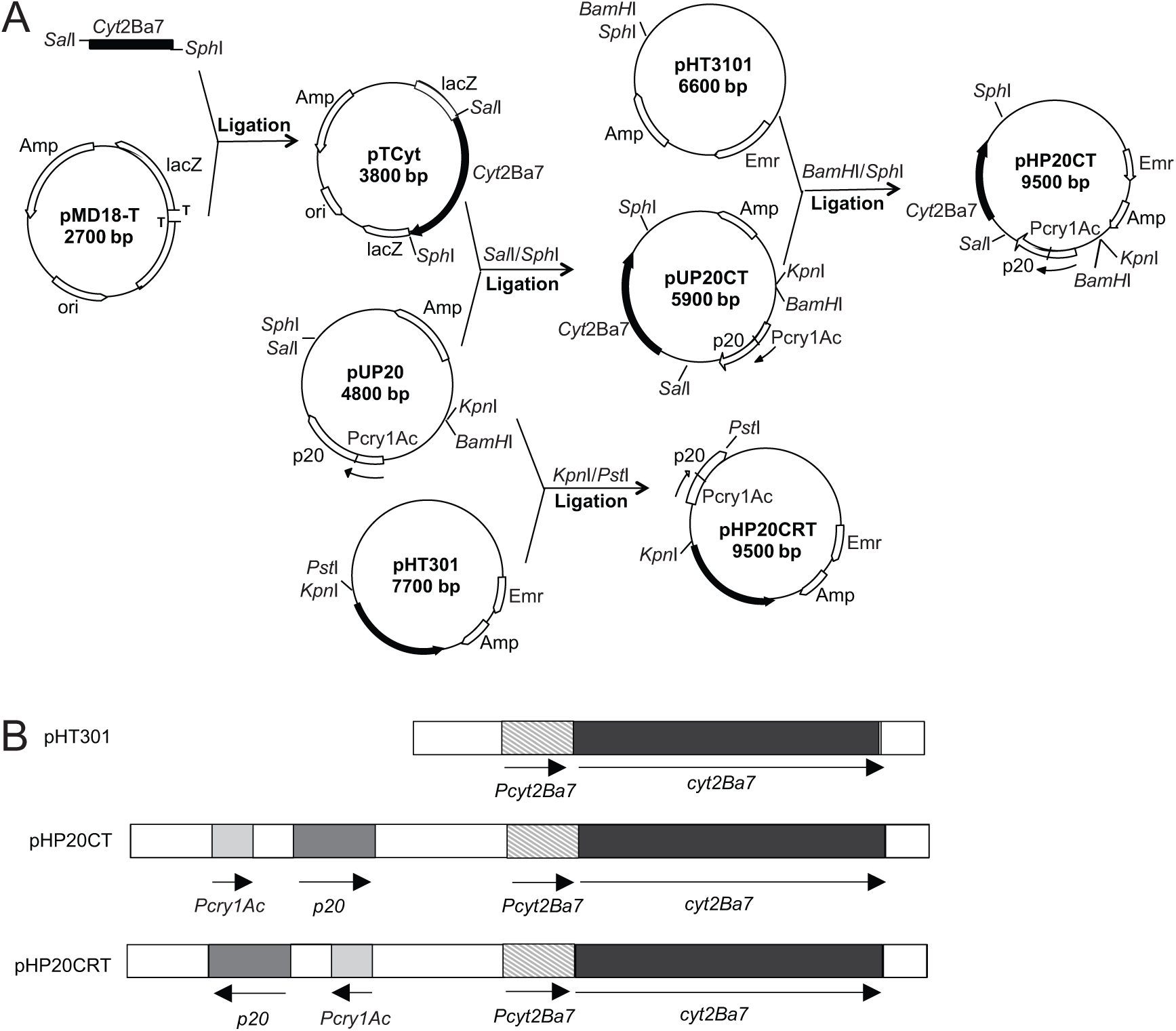
Plasmids constructing procedures (A) and maps (B) of the recombinant plasmids pHT301, pHP20CT and pHP20CRT. The plasmid pHT301 possesses *cyt*2Ba7 gene (with native promoter and ORF sequence). The plasmids pHP20CT and pHP20CRT both possess the promoter of *cry*1Ac, *p*20 gene, and *cyt*2Ba7 gene (with its native promoter) but in different directions. Arrows represent transcription direction of different genes.

### Expression of Cyt2Ba7

After inoculating 4Q7 transformed by pHT3101 (negative control), pHT301, pHP20CT and pHP20CRT in the G-Tris medium and cultured at 30°C, 1 ml medium was collected and centrifuged (15,000×g for 10 min) to harvest the cells at 12 h, 24 h, 36 h, 48 h, 60 h and 72 h after inoculation. The cell pellet was suspended in a 50 μl 1× Laemmli sample buffer containing sodium dodelyl sulfate (SDS) and boiled for 10 min, and then protein content was subjected to 12% SDS-PAGE for analysis [Shi et al., 2006]. After separation, proteins were electroblotted onto the nitrocellulose membrane (Bio-Rad, Shanghai, China) by semi-dry blotting. Antiserum for Cyt2Ba7 was used in immuoblotting analysis [Yu et al., 2002b]. The expression levels of Cyt2Ba7 in 4Q7 (pHT301), 4Q7 (pHP20CT) and 4Q7 (pHP20CRT) were analyzed from 12 to 72 h to determine the highest expression time [Shi et al., 2006].

To compare the expression levels of Cyt2Ba7 in 4Q7 (pHT301), 4Q7 (pHP20CT) and 4Q7 (pHP20CRT), these three strains were grown in G-Tris medium at 30°C for 72 hours and harvested by centrifugation as above. The cell pellets were resuspended and calibrated to the same optical densities (OD600) and 100 μl of each sample was incubated with 100 μl 2× Laemmli sample buffer and boiled. 50 μl of each sample was loaded to SDS-PAGE, stained with Coomassie Brilliant Blue R-250 and photographed by using an Electrophoresis Documentation and Analysis 120 System (Eastman Kodak Co.). The protein bands were scanned and quantified using an ImageMaster TotalLab imagining system V3 (Amersham Pharmacia Biotech) [Shi et al., 2006]. At least three replicate were performed to statistically compare Cyt2Ba7 yields.

### Transmission electron microscopy

*B. thuringiensis* and inclusion samples were purified and treated as previously described and then observed produced crystals under transmission electron microscopy JEM100cx-II (JEOL, Japan) [Nisnevitch et al., 2006; Park et al., 1998; Yu et al., 2002a].

### Immunoblotting and ligand blot analysis

To study if there is a protein-protein interaction between P20 and Cyt2Ba7, fusion proteins Cyt2Ba7 and P20 were purified as described before [Tang, 2003; Yu et al., 2002b]. One microgram purified Cyt2Ba7 was analyzed by 12% SDS-PAGE gel (Fig. 4A) and then transferred electrophoretically to a nitrocellulose membrane. To visualize immunoreactive proteins, one membrane was allowed to react directly with P20-specific antiserum [1:1000] [Tang, 2003]. Another membrane with same preparation was allowed to react with 50 μg purified P20 first and then anti-P20 antiserum [1:1000]. After washing, both membranes were incubated with secondary antibody coupled to calf intestinal alkaline phosphatase (Ap) for developing.

## Results and Discussion

### Expression of Cyt2Ba7

The plasmid pHT301 possesses both native promoter and ORF sequences of *cyt*2Ba7 (Fig. 1B). The plasmids pHP20CT and pHP20CRT both possess the native promoter of *cry*1Ac, *p*20 gene, and full-length *cyt*2Ba7 gene (with native promoter) but in different directions (Fig. 1B).

Firstly we performed a time course study on the expression of Cyt2Ba7 in *B. thruingiensis* strain 4Q7 transoformed with plasmids pHT301, pHP20CT and pHP20CRT (Fig. 2A, B and C) from 12 h to 72 h after inoculation. The immunoblot results showed that in all these three constructs, the expression of Cyt2Ba7 started from 24 h, reached the highest yield at 36 h and kept constant to 72 hours (Fig. 2). A weak band with molecular weight at around 58 kDa was observed in 4Q7 (pHP20CRT) but not in the other two constructs (Fig. 2C), which may be a dimer formation of Cyt2Ba7 [Cohen et al., 2008] and only be produced when the protein amount is high. There is no expression of Cyt2Ba7 in negative control, 4Q7 (pHT3101) (Fig. 2). Then we compared the expression of Cyt2Ba7 in 4Q7 transformed with pHT301, pHP20CT and pHP20CRT at 72 h after inoculation. Using the integrated optical density (IOD) values, we observed that the expression level of Cyt2Ba7 in 4Q7 (pHP20CRT) was increased nearly 3.2 fold than that of 4Q7 (pHT301). Cyt2Ba7 in 4Q7 (pHP20CT) was increased nearly 0.7 fold than that of 4Q7 (pHT301) (Fig. 2B). This result suggests that when *p*20 was inserted into the upstream of *cyt*2Ba7 gene, no matter which direction, the expression of Cyt2Ba7 can be significantly increased. It is very interesting that the expression of Cyt2Ba7 in 4Q7 (pHP20CRT) is much higher than that of 4Q7 (pHP20CT). The only difference between these two constructs is the direction of inserted *p*20 gene and *cry*1Ac promoter. In pHP20CT, *p*20 gene and *cry*1Ac promoter were inserted in a forward direction while in pHP20CRT, *p*20 gene and *cry*1Ac promoter were inserted in a reverse direction (Fig. 1B). Why reversely inserted *cry*1Ac promoter and *p*20 (pHP20CRT) increased the productivity of Cyt2Ba7 more significantly than forwardly inserted (pHP20CT) is unknown.

**Fig. 2.**
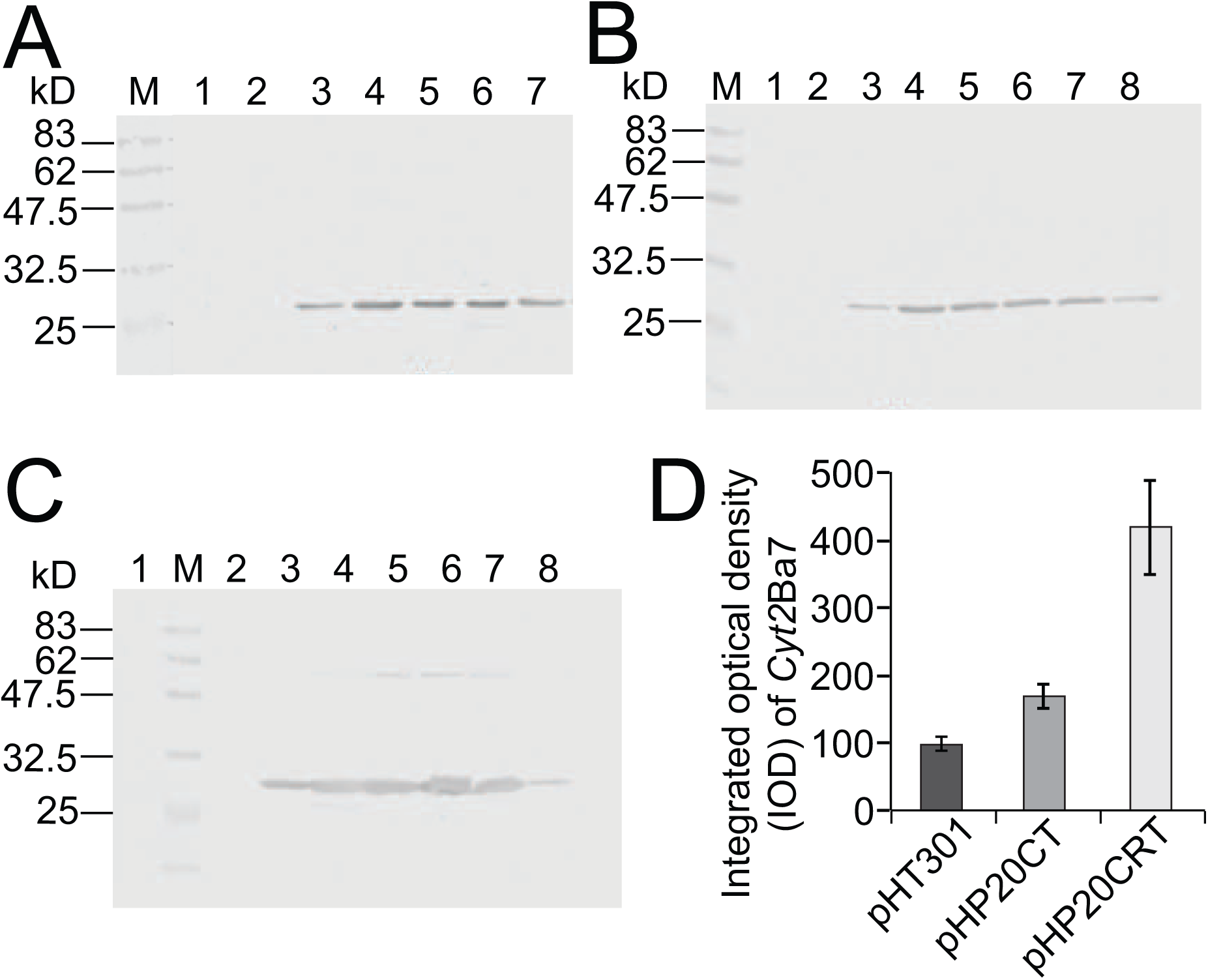
Expression time courses study of Cyt2Ba7 protein in 4Q7 transformed with plasmids pHT301(A), pHP20CT (B), pHP20CRT (C) and the expression comparison (D) of Cyt2Ba7 protein in these three strains at 72 h by using Cyt2Ba7 specific antiserum. M, protein marker; 1, 1 ml cell pellets of 72 h of 4Q7 (pHT3101); 2-7 are 1 ml cell pellet collected from 12 h, 24 h; 36 h, 48 h; 60 h and 72 h respectively. In (B) and (C), lane 8 is cell pellets of 72 h of 4Q7 (pHT301).

### The crystals of Cyt2Ba7

Then we compared the 4Q7 transformed with pHT301, pHP20CT and pHP20CRT under electron microscopy. As anticipated, we did not see any crystals from negative control, 4Q7 (pHT3101) (Fig. 3A). In 4Q7 (pHT301), we observed very few and tiny crystals of Cyt2Ba7 (Fig. 3B). Similarly, a few small irregular crystals were also observed in 4Q7 (pHP20CT) (Fig. 3C). However, from 4Q7 (pHP20CRT), we observed more and bigger hexagonal-shaped crystals ranged from 0.6 μm to 1μm (Fig. 3D, E and F). Similar results were reported on other ICPs before [Diaz-Mendoza et al., 2012; Nisnevitch et al., 2006; Park et al., 1998; Rang et al., 1996; Shao et al., 2001; Shao and Yu, 2004; Wang et al., 1997; Wu and Federici, 1993] and *Bacillus sphaericus* Bin toxin [Park et al., 2007] in *B. thuringiensis*. When *cry*1Ac promoter and *p*20 were reversely inserted in the upstream of *cyt*2Ba7, the amount and size of Cyt2Ba7 crystals are both increased, suggesting the expression of P20 can help synthesize the crystals of Cyt2Ba7.

**Fig. 3.**
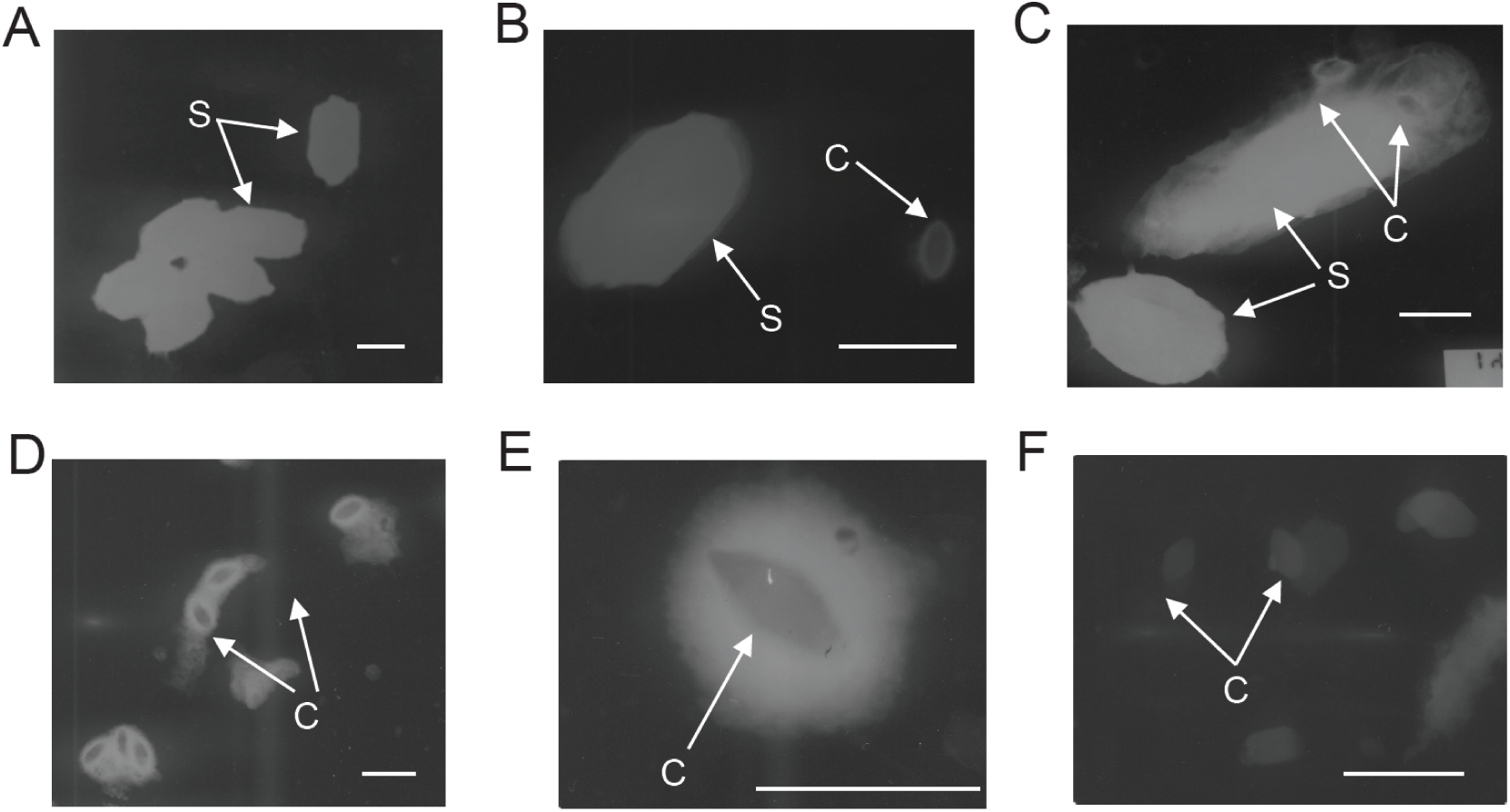
Transmission electron microscopies of crystals of Cyt2Ba7. Arrow indicates the crystals formed by Cyt2Ba7. (A) 4Q7 (pHT3101); (B) 4Q7 (pHT301); (C) 4Q7 (pHP20CT); (D) 4Q7 (pHP20CRT); (E) and (F), crystals of Cyt2Ba7 in 4Q7 (pHP20CRT). “C” stands for crystals of Cyt2Ba7 while “S” stands for spores of *B. thuringiensis*. The bar stands for 1 μm.

### The interactions between Cyt2Ba7 and P20

The molecular mechanism how P20 helps increase the synthesis of Cyt2Ba7 and other ICPs is still unknown, which can be achieved at the transcriptional, translational and posttranslational levels [Adams et al., 1989; Diaz-Mendoza et al., 2012; Park et al., 2007; Wu and Federici, 1993]. Here we insert *p*20 gene with two opposite directions into the upstream of *cyt*2Ba7 gene and the results showed that the expression level of Cyt2Ba7 in reverse form (pHP20CRT) is much higher than that of forward form (pHP20CT). The crystals synthesized are also significantly increased in amount and size. Therefore we hypothesized that P20 plays a role at protein level. For example, in the protein folding to help Cyt2Ba7 form crystals and thus prevent proteolytic degradation. If this is the case, a protein-protein interaction between P20 and Cyt2Ba7 is required and essential. To investigate this hypothesis, we used previously purified P20 proteins from *E. coli* and prepared P20-specific antiserum [Tang, 2003]. Immunoblotting and ligand blot analysis results showed Cyt2Ba7 could not be detected by P20-specific antiserum directly (Fig. 4C). However, when P20 protein was added, Cyt2Ba7 can be detected by the P20-specific antiserum (Fig. 4B). These results suggest there is a protein-protein interaction between P20 and Cyt2Ba7.

**Fig. 4.**
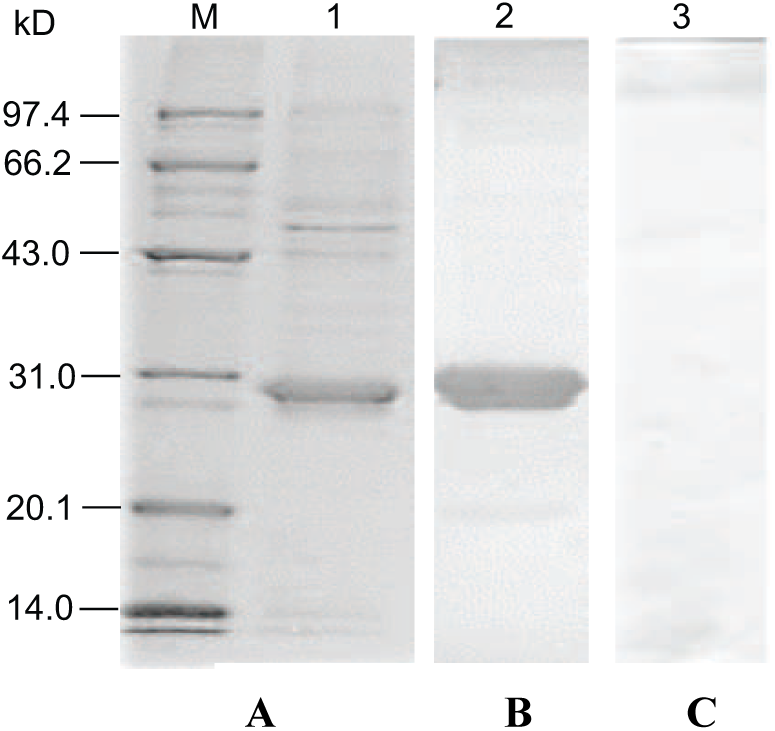
Immunoblot and ligand blot analysis of Cyt2Ba7. (A) SDS-PAGE analysis of 1 μg purified Cyt2Ba7. (B) Immunoblot analysis of Cyt2Ba7 after incubated with 50 μg purified fusion P20 protein and then P20-specific antiserum. (C) Cyt2Ba7incubated with P20-specific antiserum but without purified P20 protein. M: Standard protein marker.

P20 has been reported to enhance the expression of many other ICP proteins from *B. thuringiensis*. For example, in the presence of P20, Cyt1Aa is highly expressed and formed large ovoidal, lemon-shaped crystals [Wu and Federici, 1993]. P20 also help improve the yields of Cry1Ac [Shao et al., 2001], truncated Cry1C [Rang et al., 1996], Cry2A [Shao and Yu, 2004], Cry3A [Diaz-Mendoza et al., 2012], Cry4A [Wang et al., 1997], Cry4B [Elleuch et al., 2015] Cry11A [Park et al., 1998], Cyt2Ba [Cohen et al., 2008; Nisnevitch et al., 2006] and *B. sphaericus* Bin toxin [Park et al., 2007], which showed no or low homologies at amino acid sequences and very diverse structures. How P20 could increase the synthesis of these various proteins is still not clear. Interestingly, P20 could not promote the production of all ICP proteins. For example, P20 did not improve expression of Cry20Aa, a mosquitocidal protein [Lee and Gill, 1997]. In this study, we also found that there are certain bands on the top of Cyt2Ba7 sample (Fig. 4A), but they could not be detected by P20 and P20- specific antiserum (Fig. 4B), suggesting P20 has specificity and selectivity to bind the targeted proteins.

In summary, our study showed that P20 functions as a molecular chaperone to increase yields and facilitates crystal formation of Cyt2Ba7. Therefore it is promising for P20 to be used to increase the toxicity and mortality of *B. thuringiensis* strains to control insect pests. Many structures of ICPs including Cyt1Aa [Cohen et al., 2011], Cyt2Ba [Cohen et al., 2008] and Cry2Aa [Morse et al., 2001] have been reported. Therefore the structural functional and mutagenesis studies of P20 will assist us further explore the mechanism of P20. Better understanding P20 will pave the way to optimize its function and develop more effective and efficient insect control strategies.

## Acknowledgements

This project was supported by National Natural Sciences Foundation of China (No.39900092). Dr Wei Xu is the recipient of an Australian Research Council Discovery Early Career Researcher Award (DECRA) (DE160100382).

